# SEMA4C is a novel target to limit osteosarcoma growth, progression, and metastasis

**DOI:** 10.1101/520452

**Authors:** Branden A. Smeester, Nicholas J. Slipek, Emily J. Pomeroy, Heather E. Bomberger, Ghaidan A. Shamsan, Joseph J. Peterson, Margaret R. Crosby, Garrett M. Draper, Kelsie L. Becklin, Eric P. Rahrmann, James B. McCarthy, David J. Odde, David K. Wood, David A. Largaespada, Branden S. Moriarity

## Abstract

Semaphorins, specifically type IV, are important regulators of axonal guidance and have been increasingly implicated in poor prognoses in a number of different solid cancers. In conjunction with their cognate PLXNB family receptors, type IV members have been increasingly shown to mediate oncogenic functions necessary for tumor development and malignant spread. In this study, we investigated the role of semaphorin 4C (SEMA4C) in osteosarcoma growth, progression, and metastasis. We investigated the expression and localization of SEMA4C in primary osteosarcoma patient tissues and its tumorigenic functions in these malignancies. We demonstrate that overexpression of SEMA4C promotes properties of cellular transformation, while RNAi knockdown of SEMA4C promotes adhesion and reduces cellular proliferation, colony formation, migration, wound healing, tumor growth, and lung metastasis. These phenotypic changes were accompanied by reductions in activated AKT signaling, G1 cell cycle delay, and decreases in expression of mesenchymal marker genes *SNAI1*, *SNAI2,* and *TWIST1*. Lastly, monoclonal antibody blockade of SEMA4C *in vitro* mirrored that of the genetic studies. Together, our results indicate a multi-dimensional oncogenic role for SEMA4C in metastatic osteosarcoma and more importantly that SEMA4C has actionable clinical potential.

## 1. Introduction

Osteosarcoma is a malignant, primarily pediatric tumor, of the growing long bones with peak incidence in the second decade of life [1]. Derived from mesenchymal origins in the pre-osteoblastic lineage, osteosarcomas arise from the failure of osteoblasts to differentiate into mature bone-building cells [2]. Osteosarcomas are commonly typified by their heterogeneity, genomic instability, and frequency of systemic metastasis primarily to the lungs [3, 4]. Despite advances in chemotherapy regimens and surgical resection, survival rates for patients with osteosarcoma have remained stagnant for more than four decades [5]. The complex nature of osteosarcoma presents unique difficulties with respect to elucidating novel therapeutic targets and identifying treatment strategies that may prove most effective, particularly across individual patients. Given this, it is critically important to better understand not only the mechanisms specific to osteosarcoma development and progression, but most importantly metastasis, in order to develop better treatment options for patients with this devastating disease.

Semaphorins are a family of membrane-bound and soluble proteins that modulate a whole host of cellular functions including differentiation, cytoskeletal rearrangement, and motility [6]. Interestingly, semaphorin family members have been reported to mediate many hallmarks of cancer including cellular proliferation, angiogenesis, and immune escape [7–9]. Recent evidence from studies of the SEMA4-PLXNB family of axonal guidance molecules in normal bone cells suggests that osteoclastic expression of SEMA4D inhibits osteoblastic bone formation through suppression of IGF1 signaling [6, 10], however, recent data from our lab suggests that high expression of SEMA4D is oncogenic in osteosarcoma [11]. Given osteosarcomas retain mesenchymal-like characteristics [12], this suggests the possibility of SEMA4 members signaling through similar pathways during osteosarcomagenesis as they do during normal bone development.

Furthermore, activation of downstream signaling processes that involve the MAP kinase/PI3K pathways through heterodimerization with other receptor tyrosine kinases (RTKs) such as MET (MET proto-oncogene, receptor tyrosine kinase) and/or ERBB2 (erb-b2 receptor tyrosine kinase 2) can potentiate many of the invasive cellular processes associated with solid cancers [13]. SEMA4C, a type IV semaphorin, and its cognate receptor PLXNB2, have been recently characterized as oncogenic signaling partners in invasive breast cancer [14], hepatocellular carcinoma [15], and glioma [16]. Moreover, SEMA4C has been shown to play diverse roles in the propagation of pain signaling [17], as well as in the immune system during Th2-driven immune responses similar to other semaphorin IV family members [18].

Here, we studied SEMA4C’s role in osteosarcoma growth and metastasis. We show that 1) SEMA4C is upregulated in a subset of osteosarcoma patient samples and cell lines; 2) high SEMA4C expression enhances properties of cellular transformation, mesenchymal marker expression, and that genetic knockdown conversely reduces those phenotypes/markers; 3) SEMA4C modulates osteosarcoma growth and lung metastasis and 4) targeted monoclonal antibody blockade of SEMA4C robustly reduces these phenotypes associated with high-level SEMA4C expression. Together, this study expands upon the current known functions of SEMA4C in a highly malignant pediatric solid cancer and suggests that antibody blockade of SEMA4C-PLXNB2 signaling may overcome the current hurdles for targeting pathways that ultimately lead to metastatic lung nodule formation and continued disease progression.

## 2. Materials and Methods

### 2.1 RNA sequencing data sets

*SEMA4C* expression levels in normal human osteoblasts and human osteosarcoma patient samples were analyzed from an existing data set [11].

### 2.2 Tissue microarray (TMA) samples and scoring

The osteosarcoma tissue microarray was purchased from Biomax (Osteosarcoma: OS804c) containing 40 samples in duplicate. Slides containing 4 μm thick formalin-fixed, paraffin-embedded sections of tumor tissue were deparaffinized and rehydrated. Antigen retrieval was performed in a steamer using 1 mM Tris base EDTA buffer, pH 9.0. After endogenous peroxidase blocking, a protein block was applied. Immunohistochemistry (IHC) for SEMA4C was performed using a rabbit anti-SEMA4C primary antibody (#AF6125, R & D Systems) on an autostainer (Dako). Detection was achieved using the Envision rabbit detection system (Dako) with diaminobenzidine (DAB) as the chromogen. Tissue sections were imaged on a Nikon E800M microscope at 40X magnification using a Nikon DSRi2 camera and Nikon Elements D Version 4 software. Slides were evaluated and scored as previously described [19].

### 2.2 Cell culture

All osteosarcoma cell lines were purchased and obtained from the American Type Culture Collection (ATCC). Normal human osteoblasts (NHOs) and primary human umbilical vein cells (HUVECs) were purchased from Lonza. All cell lines were grown and maintained in accordance with standard cell culture techniques. Both HOS and MG-63 cells were grown in DMEM. G-292 cells were grown in McCoy’s 5A. SJSA-1 cells were grown in RPMI-1640. NHOs were grown in αMEM. Osteosarcoma cell line media was fortified with 10% fetal bovine serum (FBS) and 1X penicillin/streptomycin and NHO media was fortified with 20% FBS and 1X penicillin/streptomycin. HUVECs were cultured with EGM-2 bullet kit media (#cc-3162, Lonza) with 1X antibiotic-antimycotic (#15240062, Thermo Fisher) in cell culture flasks coated with 1% gelatin (#G1383, Sigma Aldrich)‥ All cell cultures were incubated in a water-jacketed incubator set at 5% carbon dioxide (CO_2_) and at 37°C. All cell lines except NHO, G-292, and HUVECs were authenticated by the University of Arizona Genetics Core (UAGC) using short tandem repeat profiling. All cell lines were found to be free from mycoplasma.

### 2.3 Generation of shRNA knockdown and overexpression cell lines

Stable knockdown of SEMA4C was accomplished with pGIPZ lentiviral vectors expressing an shRNA against SEMA4C in conjunction with GFP and a puromycin selection marker (sh1 #V3LHS_394644, sh2 #V3LHS_413698, Open Biosystems). Control pGIPZ vector with non-targeting shRNA was used as a control (#RHS4346, Open Biosystems). Lentiviral particles were produced with 293T cells co-transfected with pGIPZ shRNAs, pMD2.G envelope (#12259, addgene) and psPAX2 (#12260, addgene) packaging vectors. Stable shRNA knockdown lines were established via puromycin selection at 1 μg/mL. Overexpression vectors were generated using the human *SEMA4C* cDNA sequence (#40035253, Dharmacon) and vectors/cell lines were established using previously described methodology [20, 21].

### 2.4 Anti-SEMA4C antibody treatment *in vitro*

Anti-SEMA4C or isotype control antibody was administered at 10 μg/mL and 15 μg/mL where indicated (#sc-136445, Santa Cruz Biotechnology).

### 2.5 RNA isolation and Quantitative RT-PCR

Total RNA was extracted from cell lines using the High Pure RNA Isolation Kit (Roche, Basel). 1 μg of extracted RNA was reverse transcribed into cDNA using the Transcriptor First Strand Synthesis kit (Roche). Quantitative RT-PCR was performed in triplicate using SYBR green mix (Qiagen) on an ABI 7500 machine (Applied Bio Systems). Primer sequences are available in Suppl. Table 2. All measurements were calculated using the ΔΔCT method and expressed as fold change relative to respective control non-silencing shRNA line (shCON).

### 2.6 ELISA

Quantification of levels of soluble SEMA4C secreted in the conditioned media (72 hours) of indicated cell lines was performed via manufacturer’s instructions (#MBS705730, MyBioSource).

### 2.7 Western blot

Protein was extracted from cultured cells in RIPA buffer containing a protease inhibitor (Roche) and phosphatase inhibitors (Sigma-Aldrich). Total protein was quantified using BCA (Thermo Fisher). Cell lysate was loaded and run on 4-12% Bis-Tris gels (Thermo Fisher) and transferred to PVDF membranes (Bio-Rad). Membranes were blocked in 5% nonfat dry milk for 1 hour (Bio-Rad) and incubated gently shaking overnight at 4°C in 1° antibody/PBS-T. Subsequently, membranes were washed and incubated in conjugated 2° antibodies for 1 hour. Blots were thoroughly washed following 2° incubation and developed using the WesternBright Quantum detection kit (Advansta) and the LICOR Odyssey (LICOR). A complete list of antibodies and other reagents utilized is available in Supp. Table 1. Densitometry was performed on all images. Bands were compared to each respective loading control, normalized to 1, and compared.

### 2.8 MTS cellular proliferation assay

Cellular proliferation assays were performed as previously described [11]. Briefly, modified cells (1.2 × 10^3^) were plated per well in 96-well plates. Cells were measured at 24, 48, 72, and 96 hours post plating. Absorbances at 490 nm and 650 nm were read using a SynergyMx (BioTek) fluorescence plate reader.

### 2.9 Transwell migration assay

Modified cells (2.5 × 10^4^) were seeded in 500 μL of serum free media in the upper chamber of 8 μm inserts (Corning). The lower chamber was filled with 750 μL media fortified with 10% fetal bovine serum (FBS) as a chemo attractant. After 24 hours, non-migrating cells where removed with a cotton swab. Migrated cells located on the lower side of the chamber were fixed with crystal violet, air-dried, and photographed to quantify migration of cells. For anti-SEMA4C studies, cells were seeded into the top chamber with antibody following a 6-hour pre-treatment at indicated concentration.

### 2.10 Wound healing assay

Wound healing assays were performed as previously described [20]. Briefly, 1 × 10^4^ cells were plated into a removable 2-well silicone culture insert which generated a defined cell-free gap (ibidi). Cells were then incubated for 24 hours before inserts were removed and fresh cell culture media was added. Phase contrast images of the wounds were acquired every 10 minutes for 16-20 hours, which was sufficient for the cells to completely close the simulated wound gap. A custom-written image segmentation algorithm in MATLAB was used to measure wound distance over time and to calculate closure rate.

### 2.11 3D microfluidic cellular adhesion assay

Cellular adhesion assays were performed as previously described [22]. Briefly, the microfluidic model was fabricated using standard soft lithography of polydimethylsiloxane (PDMS) (#4019862, Ellsworth). Rat tail collagen I (#CD354249, Corning) was buffered with PBS and cell culture grade water to create a 6 mg/mL solution. Collagen was then loaded into the lower channel of the microfluidic model and allowed to nucleate at 37°C for at least 1 hour. The upper channel was coated with 1% gelatin solution for 30 minutes at room temperature. HUVECs were released using trypsin, resuspended in a 4% w/v dextran (#31392, Sigma Aldrich) solution in EGM-2 media at a concentration of approximately 1 × 10^6^ cells per mL, and 50 μL of the cell solution was added to the inlet of the microfluidic model. HUVECs become confluent in the channel over 24-48 hours at 37°C. 5 × 10^4^ green fluorescent protein (GFP)-expressing osteosarcoma cells were then added to the inlet of the model and allowed to adhere over 3 hours. Microfluidic models were imaged at 24 hours. Adhesion and invasion of osteosarcoma cells was quantified.

### 2.12 Soft agar colony formation assay

Modified cells (1 × 10^4^) were seeded into a 0.35% agar solution placed on top of a 0.5% agar in six-well plates and allowed to incubate for 2-3 weeks. The resultant colonies were fixed, divided into four quadrants, and imaged using microscopy. Colonies were quantified via ImageJ v1.52a software using a standard colony quantification macro [11].

### 2.13 Flow cytometry

Cells were fixed with 2% paraformaldehyde (Electron Microscopy Sciences) and permeabilized with cold 90% methanol (Sigma). Cell cycle analysis was performed using PI/RNase Staining Buffer (BD Pharmingen). Cleaved caspase 3 (Asp175, clone D3E9) PE was purchased from Cell Signaling Technologies and cells were stained according to manufacturer’s recommendations. Cells were analyzed on an LSR II or Fortessa digital flow cytometer (BD Biosciences) at the University of Minnesota Flow Cytometry Resource. Analysis was performed using FlowJo software.

### 2.14 Orthotopic osteosarcoma mouse model

All animal procedures were performed in accordance with protocols approved at the University of Minnesota in conjunction with the Institutional Animal Care and Use Committee (IACUC). Modified cells (2.5 × 10^5^) were injected into the calcaneous of 6-8-week-old immunocompromised mice (NOD Rag Gamma, Jackson Labs) [23–25]. Tumor volume was calculated via caliper measurements using the formula V = (W*W*L)/2 where V equals tumor volume, W equals tumor width and L equals tumor length [26].

### 2.15 Lung metastasis evaluation

Micrometastatic nodules were examined via H & E histology at 4X magnification. Quantification of nodule number and area was undertaken in 9 sections from four mice/group (shCON and shSEMA4C) at 4X using ImageJ v1.52a software.

### 2.16 Statistical analysis

All statistical analyses were performed using Prism v8 software (GraphPad). All data are presented as mean ± standard error of the mean (SEM). Two groups were compared using a two-tailed unpaired Student’s t-test. Three or more groups were compared using a One-way ANOVA with Bonferroni’s post hoc or Two-way ANOVA analyses were performed and followed with Bonferroni’s post hoc testing. All statistical analyses are individually indicated where applied throughout. In all cases, *p* < 0.05 was considered statistically significant.

## 3. Results

### SEMA4C is highly expressed in some osteosarcoma tissues and cell lines

In order to investigate the potential role of SEMA4C in osteosarcoma development and metastasis, we first examined *SEMA4C* mRNA expression in 12 human osteosarcoma clinical samples and in 3 normal human osteoblast control samples (Fig. 1A). Human osteosarcoma samples had significantly higher *SEMA4C* expression when compared to normal human osteoblasts, (Fig. 1A; \**p* < 0.05). Next, we examined SEMA4C protein expression in a commercially available human osteosarcoma TMA (Figs. 1B and C). When all 40 sections were quantified, 32.5% had weak expression, 47.5% had moderate expression, and 20.0% had strong expression of SEMA4C (Fig. 1B). Representative images of each staining pattern are shown in Fig. 1C. The majority of tissue sections were greater than 75% SEMA4C positive (Supp. Fig. 1A) and had primarily cytoplasmic/membraneous staining localization (Supp. Fig. 1B). These results indicate that a subset of osteosarcoma patient tissues express high levels of SEMA4C. We next probed a number of commercially available osteosarcoma cell lines for SEMA4C and cognate receptor PLXNB2 expression via western blot and compared them to normal human osteoblasts (NHO). Relative to the NHOs, most osteosarcoma cell lines examined expressed detectable membrane-bound SEMA4C (Fig. 1D). Both MG-63 and G-292 cells had no detectable expression in Fig. 1D, but was minimally detectable in Fig. 2A at high exposures. Since SEMA4C is known to be shed from the extracellular membrane [27], we also analyzed cell culture media of indicated cell lines for the presence of soluble SEMA4C (Fig. 1E). Soluble SEMA4C was detectable in all lines including NHO, however, soluble SEMA4C was expressed at significantly elevated levels in U2OS, HOS, SJSA-1, and MG-63 osteosarcoma cell lines as compared to NHOs (Fig. 1E, \**p* < 0.05, \*\**p* < 0.01). These data indicate that SEMA4C is upregulated in a subset of osteosarcoma tissues and cell lines compared to normal osteoblasts.

**Fig. 1.**
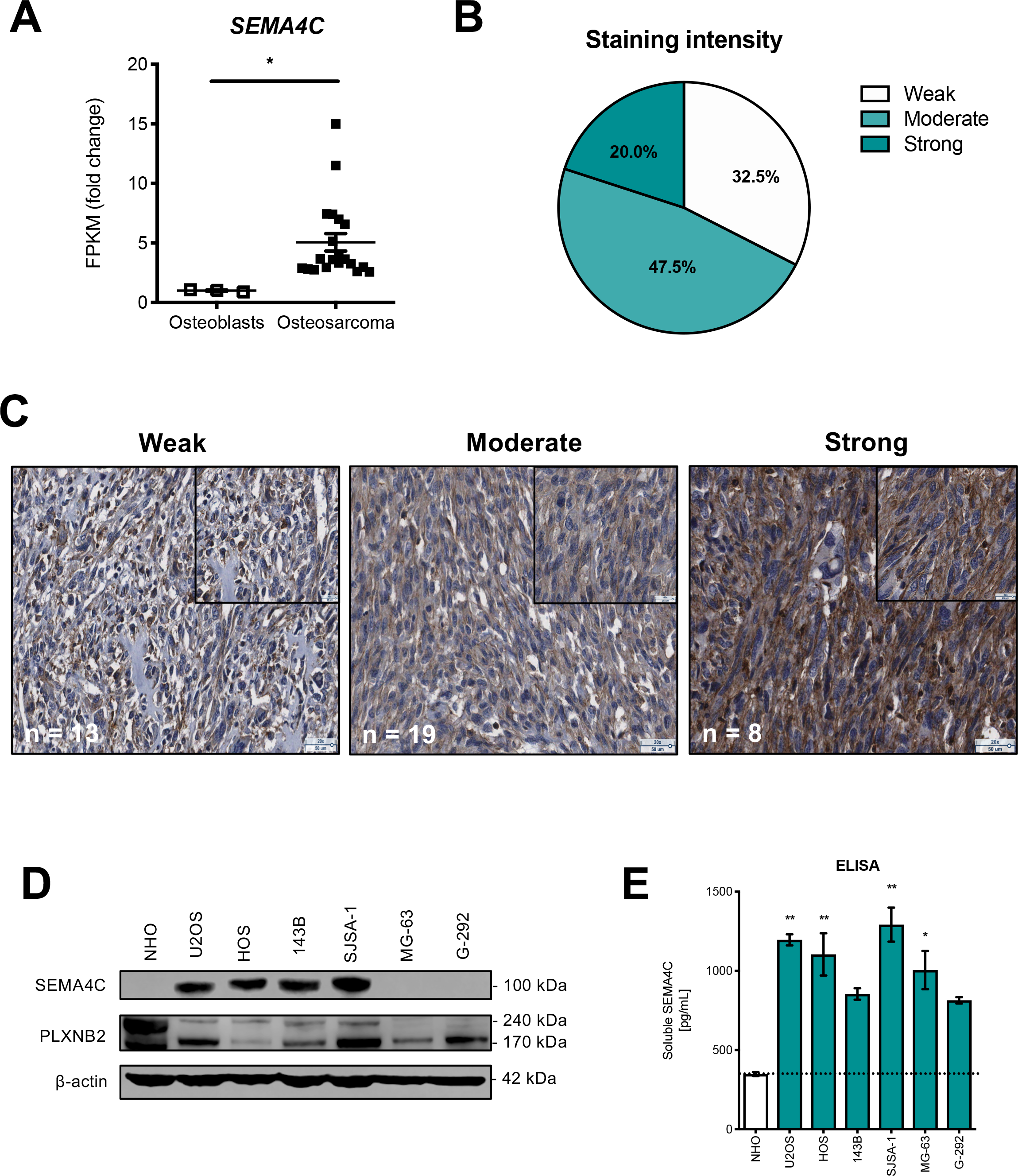
SEMA4C is upregulated in a subset of osteosarcoma tissue samples and cell lines. **A.** Relative *SEMA4C* RNA expression levels in normal human osteoblasts (n = 3) compared to osteosarcoma patient samples (n = 12). Data shown are fold change compared to osteoblasts ± SEM; \**p* < 0.05; unpaired Student’s T-test. **B.** Summary of scores identified in human osteosarcoma TMA following staining with anti-SEMA4C. **C.** Representative images of SEMA4C staining with each expression intensity and number of sections with indicated staining. Representative images are shown at 20X; insets are 40X. Scale bars = 25 and 50 μm where indicated. **D.** Western blots of SEMA4C and cognate receptor PLXNB2 expression in normal human osteoblasts (NHO) and osteosarcoma cell lines. **E.** ELISA analysis of soluble SEMA4C expression in NHOs and osteosarcoma cell supernatants. Data shown as mean ± SEM (n = 2/group); \**p* < 0.05, \*\**p* < 0.01; One-way ANOVA.

**Fig. 2.**
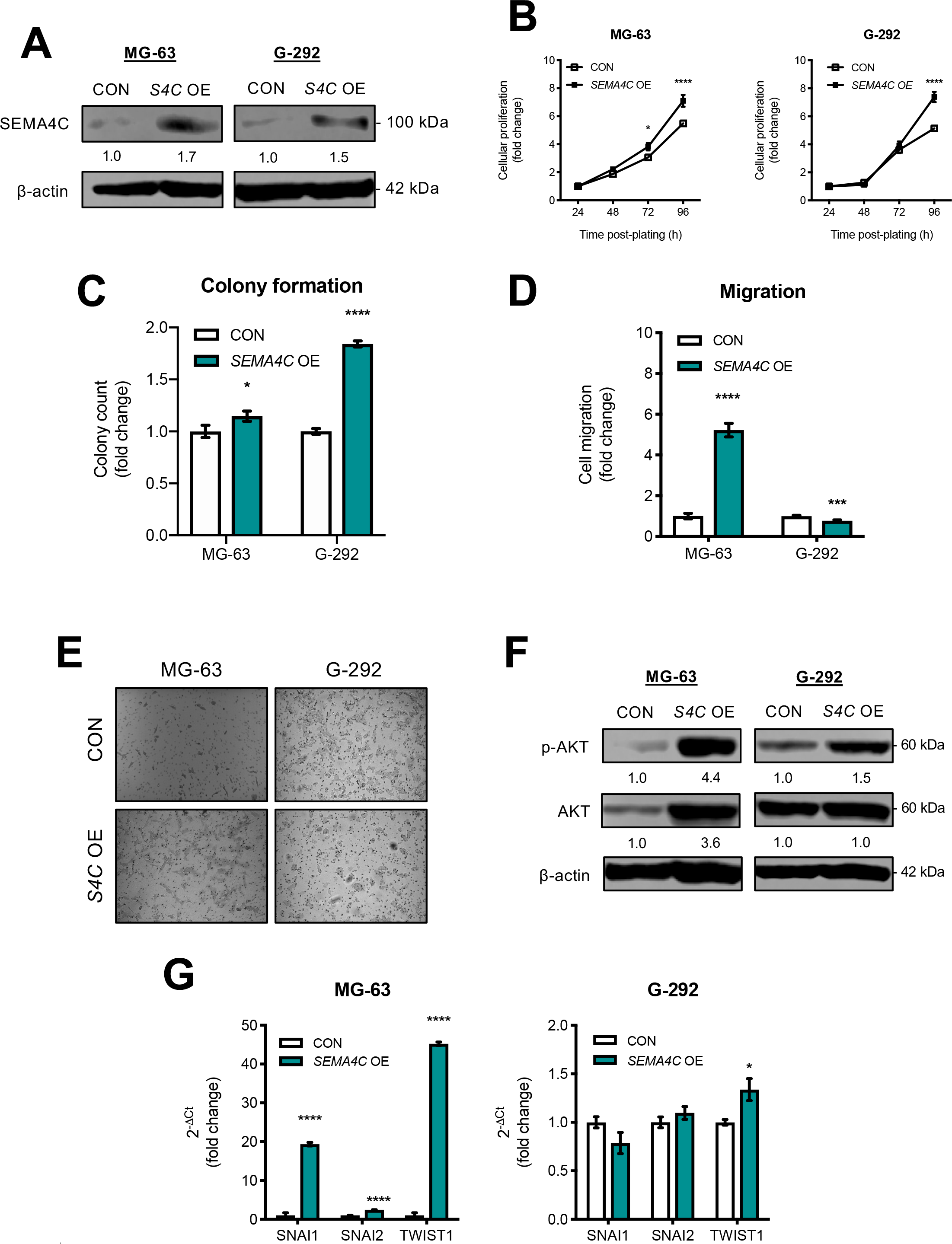
*SEMA4C* overexpression promotes increased cellular growth, colony formation, and migration in osteosarcoma cell lines. **A.** Western blots confirming overexpression of SEMA4C. **B.** *SEMA4C* overexpression increases the proliferation of osteosarcoma cell lines. Data shown as fold change ± SEM (n = 18/group); \**p* < 0.05, \*\*\*\**p* < 0.0001; Two-way ANOVA. **C.** *SEMA4C* overexpression promotes anchorage-independent growth. Data as fold change ± SEM (n = 36/group); \**p* < 0.05, \*\*\*\**p* < 0.0001; unpaired Student’s T-tests. **D.** *SEMA4C* overexpression modulates migration. Data as fold change ± SEM (n = 12/group); \*\*\**p* < 0.001, \*\*\*\**p* < 0.0001; unpaired Student’s T-tests. **E.** Representative images of cellular migration. **F.** Western blots of activated AKT signaling in *SEMA4C*-overexpressing cell lines. **G.** Relative expression of mesenchymal marker genes in cell lines ± *SEMA4C* overexpression (n = 3/group); multiple Student’s T-tests.

### Overexpression of *SEMA4C* promotes facets of cellular transformation in osteosarcoma cells

To understand the SEMA4C-PLXNB2 signaling axis in osteosarcoma, we overexpressed *SEMA4C* in endogenous low expressing MG-63 and G-292 osteosarcoma cell lines and confirmed by western blot (Fig. 2A). Overexpression of *SEMA4C* increased proliferation in both lines (Fig. 2B, \**p* < 0.05, \*\*\*\**p* < 0.0001). In a soft agar colony formation assay, overexpression of *SEMA4C* promoted modest increases in colony formation in both lines (Fig. 2C, \**p* < 0.05, \*\*\*\**p* < 0.0001). Interestingly, in a transwell migration assay, *SEMA4C*-overexpressing MG-63 cells had increased migration while G-292 cells displayed slightly reduced migration (Fig. 2D, \*\*\**p* < 0.001, \*\*\*\**p* < 0.0001). Representative images of the migration experiments are shown in Fig. 2E. Both MAPK and PI3K signaling were investigated following phenotypic assays. These changes in proliferation, migration, and colony formation were associated with increased activation of AKT signaling, but not ERK signaling (data not shown) (Fig. 2F). Lastly, overexpression of *SEMA4C* significantly promoted upregulation of mesenchymal markers *SNAI1*, *SNAI2*, and *TWIST1* in MG-63 cells, while all but *TWIST1* remained largely unchanged in G-292 (Fig. 2G, \**p* < 0.05, \*\*\*\**p* < 0.0001). These data suggest that overexpression of SEMA4C promotes facets of cellular transformation and mesenchymal marker expression.

### Knockdown of SEMA4C reduces cellular proliferation and colony formation

To complement our gain-of-function (GOF) studies and to elucidate the effects of knockdown of SEMA4C on cellular proliferation and anchorage-independent growth, we performed loss-of-function (LOF) experiments in two endogenously high SEMA4C-expressing lines (See Figs. 1D and 1E). HOS and SJSA-1 osteosarcoma cell lines were transduced with shRNAs against SEMA4C (shSEMA4C or shS4C abbreviated) or non-silencing control (shCON) and stably selected with puromycin. Confirmation of optimal shRNA knockdown was evaluated via qRT-PCR (Fig. 3A, \*\**p* < 0.01, \*\*\*\**p* < 0.0001) and western blot (Fig. 3B). Following evaluation, shRNA #2 was chosen for both lines in all subsequent experiments. Knockdown of SEMA4C reduced cellular proliferation (Fig. 3C, \*\*\*\**p* < 0.0001) and colony formation (Fig. 3D, \*\*\*\**p* < 0.0001) in both lines. These reductions in 2D and 3D growth were accompanied by downregulation of activated AKT signaling (Fig. 3E). Lastly, silencing of SEMA4C was associated with G1 cell cycle delay in both lines (Fig. 3F, \**p* < 0.05, \*\**p* < 0.01). Silencing of SEMA4C did not induce cleaved CASP3 activity (Supp. Figs. 2A and 2B). Together, these data suggest SEMA4C modulates cellular growth and colony formation in osteosarcoma cell lines.

**Fig. 3.**
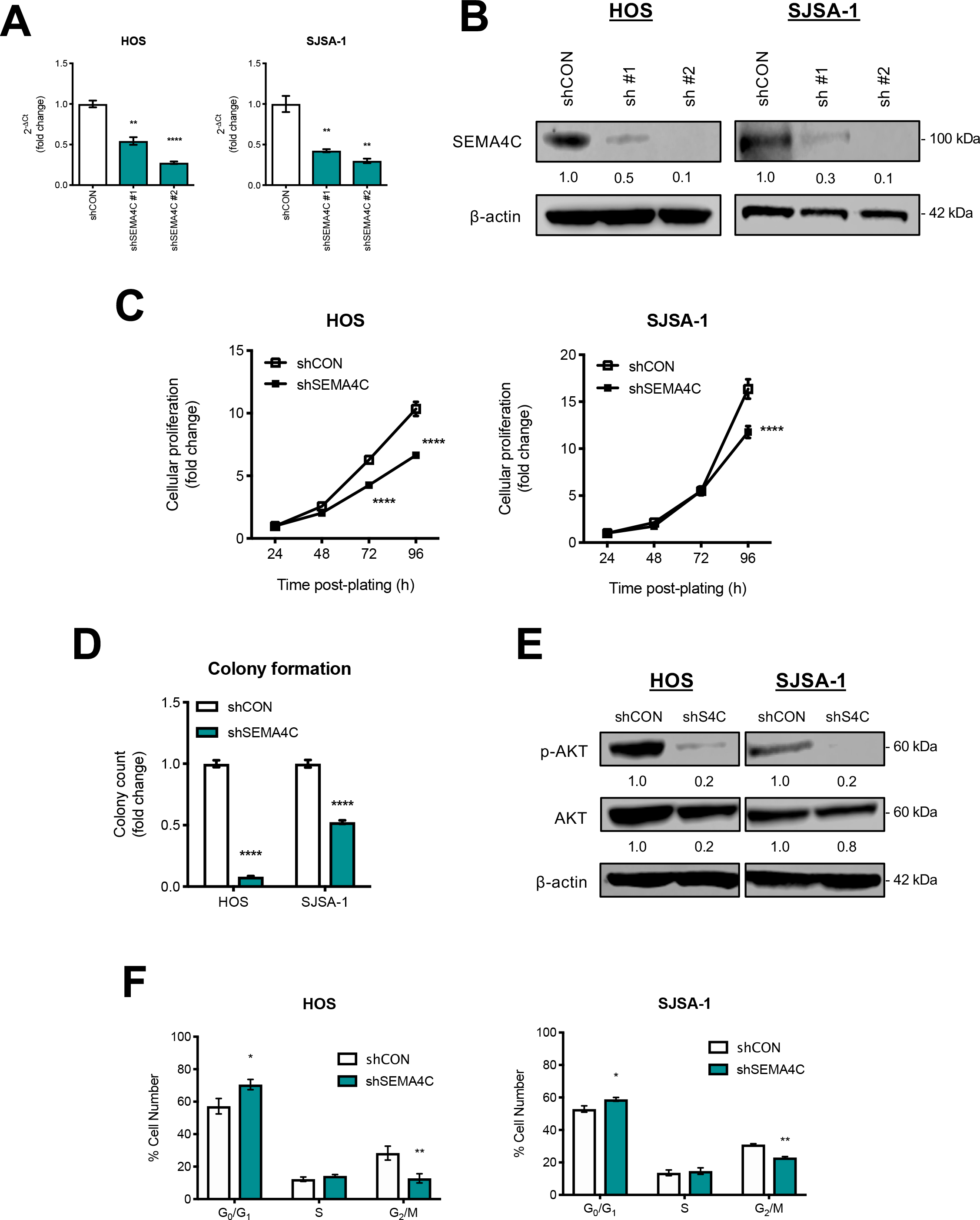
Knockdown of SEMA4C reduces cellular growth, colony formation, and promotes cell cycle delay. **A.** Confirmation of SEMA4C knockdown via qRT-PCR. Data shown as fold change ± SEM (n = 3/group); \*\**p* < 0.01, \*\*\*\**p* < 0.0001; One-way ANOVA. **B.** Western blots of SEMA4C in control shCON and shSEMA4C cell lines. **C.** Knockdown of SEMA4C reduces cellular proliferation. Data as fold change ± SEM (n = 18/group); \*\*\*\**p* < 0.0001; Two-way ANOVA. **D.** Colony formation in SEMA4C knockdown cell lines. Data as fold change ± SEM (n = 36/group); \*\*\*\**p* < 0.0001; unpaired Student’s T-test. **E.** Western blots of activated AKT signaling in SEMA4C knockdown cells. **F**. Silencing of SEMA4C induces G1 cell cycle delay; Data shown as number of cells ± SEM (n = 3/group); Two-way ANOVA.

### SEMA4C promotes cellular motility and loss of adhesion

Next, we examined the role of SEMA4C in cellular movement using migration chambers, wound healing assays, and 3D microfluidic chambers. Knockdown shSEMA4C lines displayed reduced cellular migration (Figs. 4A, \*\**p* < 0.01, \*\*\*\**p* < 0.0001). Representative migration images are shown in Fig. 4B. Following the migration assay, we evaluated cellular adhesion and invasion using 3D microfluidic chambers (Figs. 4C and 4D and Supp Figs. 3A-D). Increased adhesion was observed in HOS, but not SJSA-1 cells (Fig. 4C, \**p* < 0.05). Representative images 24 hours post are depicted in Fig. 4D. No changes in invasion were observed in either HOS or SJSA-1 knockdown cells (data not shown). Similarly, SJSA-1 knockdown cells showed reduced wound closure rates while a non-significant trend towards reduced closure rates was observed in HOS (Fig. 4E, *p* = 0.06, \**p* < 0.05). Representative wound healing photographs are shown in Supp. Fig. 4A. These changes in cell motility were associated with reductions in expression of mesenchymal markers *SNAI1*, *SNAI2,* and *TWIST1* in both knockdown lines (Fig. 4F, \*\*\**p* < 0.001, \*\*\*\**p* < 0.0001). Together, these data suggest SEMA4C promotes cellular motility and loss of adhesion in osteosarcoma cell lines.

**Fig. 4.**
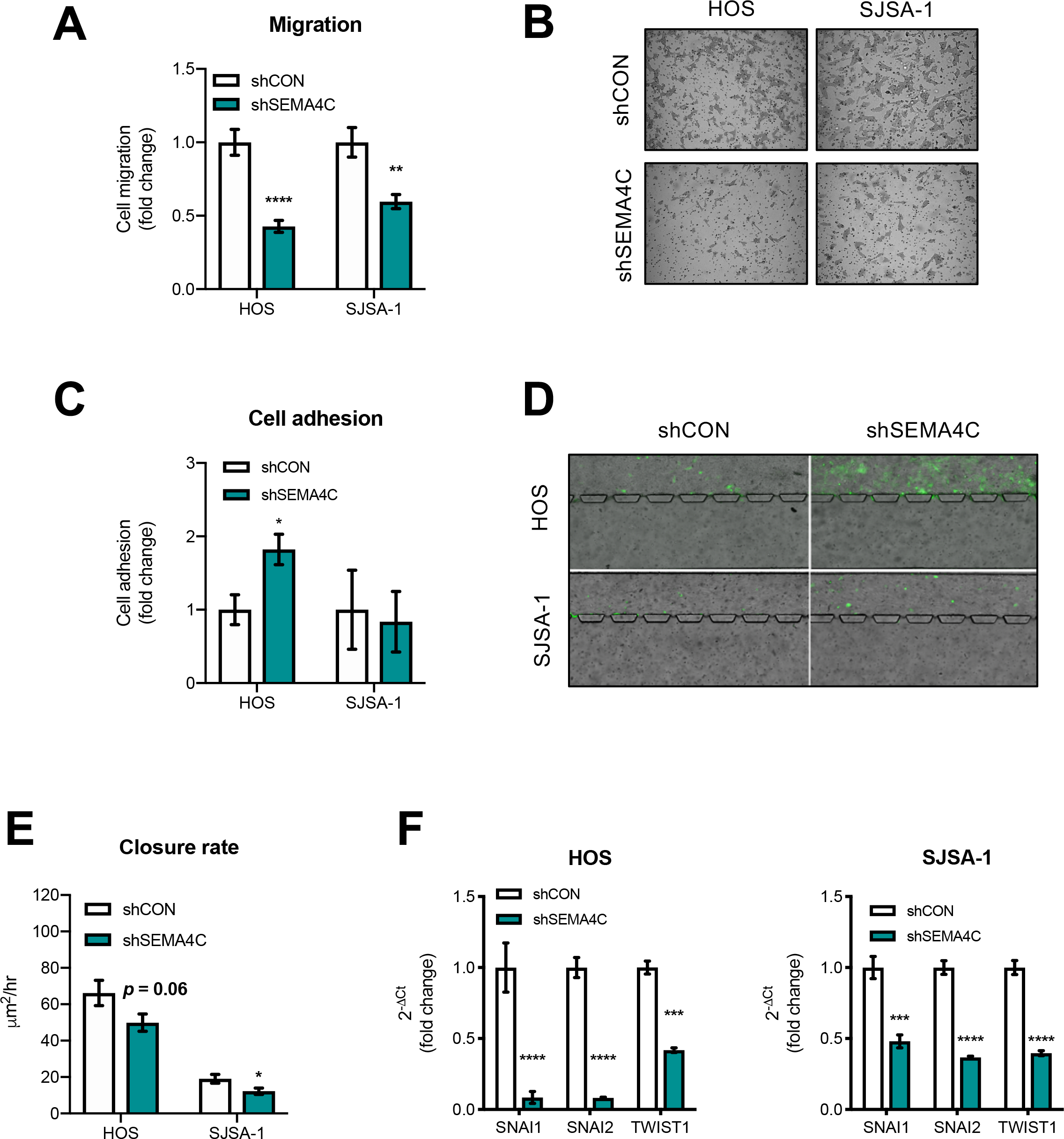
SEMA4C knockdown reduces cell motility, promotes adhesion, and downregulates mesenchymal marker expression. **A.** Knockdown of SEMA4C reduces cellular migration. Data as fold change ± SEM (n = 12/group); \*\**p* < 0.01, \*\*\*\**p* < 0.0001; unpaired Student’s T-test. **B.** Representative cellular migration images in knockdown cell lines. **C.** Silencing of SEMA4C increases cellular adhesion in HOS cells (n = 5-6/group); \**p* < 0.05; unpaired Student’s T-test. **D.** Representative images of cellular adhesion 24 hours post. **E.** Wound closure rate in SEMA4C-deficient cells (n = 36/group); \**p* < 0.05; unpaired Student’s T-test. **F.** qRT-PCR of mesenchymal markers. Data as fold change ± SEM (n = 3/group); \*\*\**p* < 0.001, \*\*\*\**p* < 0.0001; multiple Student’s T-tests.

### SEMA4C knockdown reduces tumor growth and development of lung metastases

Next, we evaluated the effects of SEMA4C knockdown on osteosarcoma tumor growth and lung metastasis in an orthotopic mouse model. Following injection of either shCON or shSEMA4C knockdown cells into the calcaneous of immunodeficient mice, tumors were allowed to form, and caliper measurements were taken beginning 10 days post-implantation and every 5 days for 30 days. Both HOS and SJSA-1 knockdown cell lines displayed reduced tumor growth (Fig. 5A, \**p* < 0.05, \*\*\**p* < 0.001, \*\*\*\**p* < 0.0001). Representative gross images of lungs from HOS shCON and shSEMA4C mice are shown in Fig. 5B. Visible macrometastatic nodules are indicated by white arrows. Whole cell lysates were made from two representative control and knockdown tumors from each cell line. SEMA4C knockdown was confirmed *in vivo* and activated AKT signaling also reduced in SEMA4C knockdown tumors (Fig. 5C). These results mirrored that of our GOF and LOF *in vitro* studies in Fig. 2F and Fig. 3F respectively. Considering metastatic capacity is often associated with increased cell growth, motility, and anchorage-independence [28, 29], we investigated the effects of SEMA4C knockdown on lung nodule formation and size. When lung sections were examined, both SEMA4C knockdown lines had reduced numbers of micrometastatic nodules (Fig. 5D, \*\*\*\**p* < 0.0001) and nodule area (Fig. 5E, \*\*\*\**p* < 0.0001). Representative lung H & E images and high powered-insets are shown in Fig. 5F for both cell lines. Black arrows indicate micrometastases. Together, SEMA4C promotes tumor growth and lung metastasis formation in osteosarcoma.

**Fig. 5.**
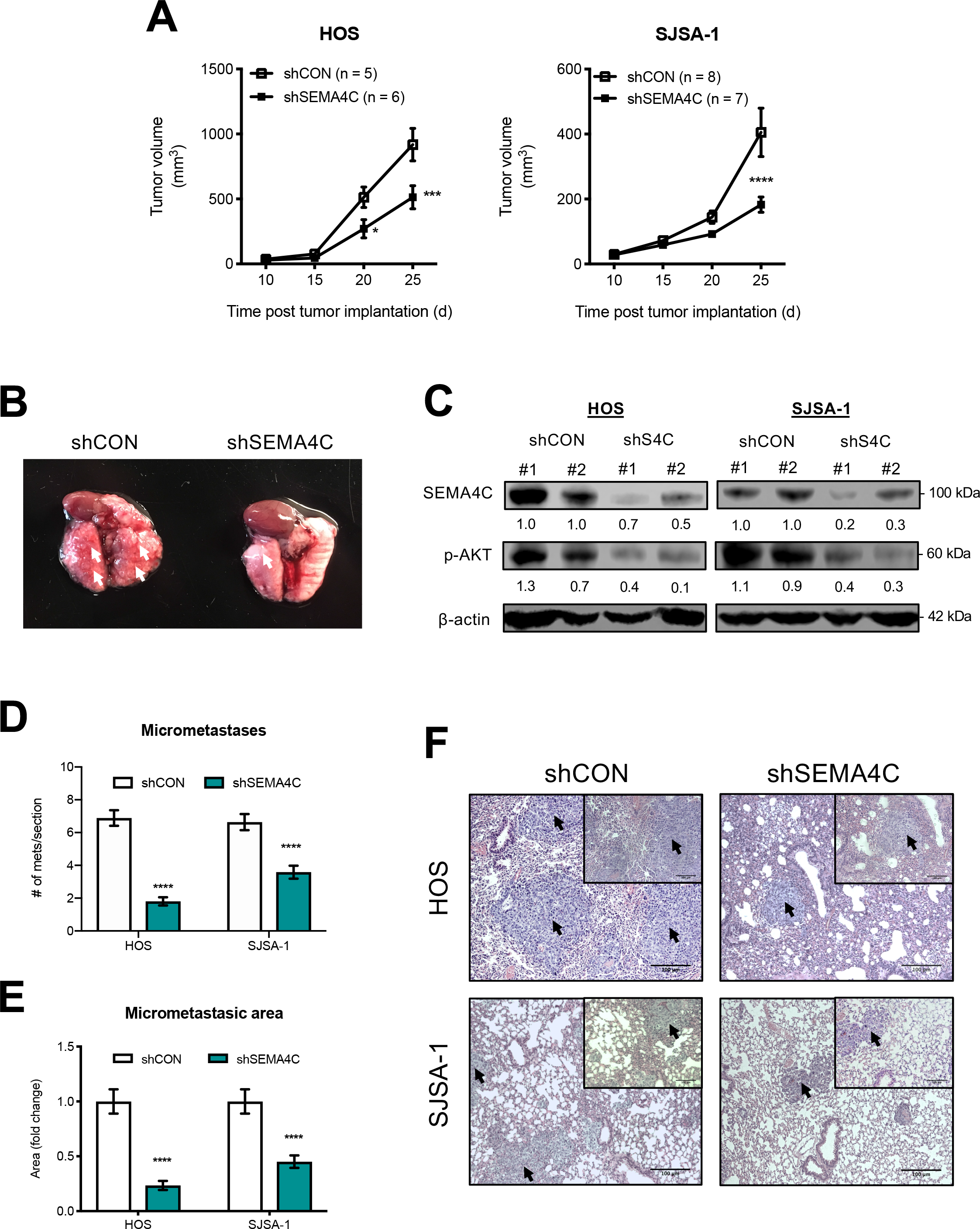
SEMA4C knockdown decreases osteosarcoma tumor growth and lung metastasis. **A.** Tumor volume measurements in SEMA4C knockdown orthotopic injections. Data shown as mean volume in mm^3^ ± SEM; \**p* < 0.05, \*\*\**p* < 0.001, \*\*\*\**p* < 0.0001; Two-way ANOVA. **B.** Representative gross lung images from a HOS control and SEMA4C knockdown animal. White arrows indicate macrometastatic nodules. **C.** Western blot images in two control and SEMA4C knockdown animals from each cell line confirming SEMA4C knockdown and reductions in AKT signaling. **D.** Number of micrometastatic lung nodules/section. Data shown as mean number of nodules ± SEM (n = 36/group); \*\*\*\**p* < 0.0001; unpaired Student’s T-test. **E.** Area measurements of micrometastases in sections. Data shown as mean area ± SEM (n = 36/group); \*\*\*\**p* < 0.0001; unpaired Student’s T-test. **F.** Representative H & E lung images are shown at 20X; insets are 40X. Black arrows indicate micrometastases, Scale bars = 100 μm.

### Monoclonal antibody blockade of SEMA4C reduces tumorigenic properties of osteosarcoma cell lines

Lastly, we sought to evaluate the therapeutic potential of SEMA4C blockade using a commercially available monoclonal antibody raised against amino acids 400-510 of human SEMA4C. To evaluate the effects of blockade *in vitro*, we treated wild-type normal human osteoblasts (NHO), HOS, and SJSA-1 osteosarcoma cells with two concentrations of anti-SEMA4C or isotype control IgG and assayed its effects on cellular proliferation. Reduced cellular proliferation was observed following a 48-hour treatment in both osteosarcoma cell lines, but not NHOs (Fig. 6A, \*\*\*\**p* < 0.0001). Similarly, treatment with anti-SEMA4C also reduced migration in both osteosarcoma cell lines (Fig. 6B, \*\*\*\**p* < 0.0001). Representative images are shown in Fig. 6C. Following treatment, G1 cell cycle delay was again observed similar to our genetic studies in both cell lines (Fig. 6D, \**p* < 0.05, \*\**p* < 0.01, \*\*\**p* < 0.001, \*\*\*\**p* < 0.0001). Our findings suggest that anti-SEMAC treatment may prove valuable for combating osteosarcoma tumor growth, progression, and ultimately lung metastasis through disruption of oncogenic SEMA4C signaling (Fig. 6E).

**Fig. 6.**
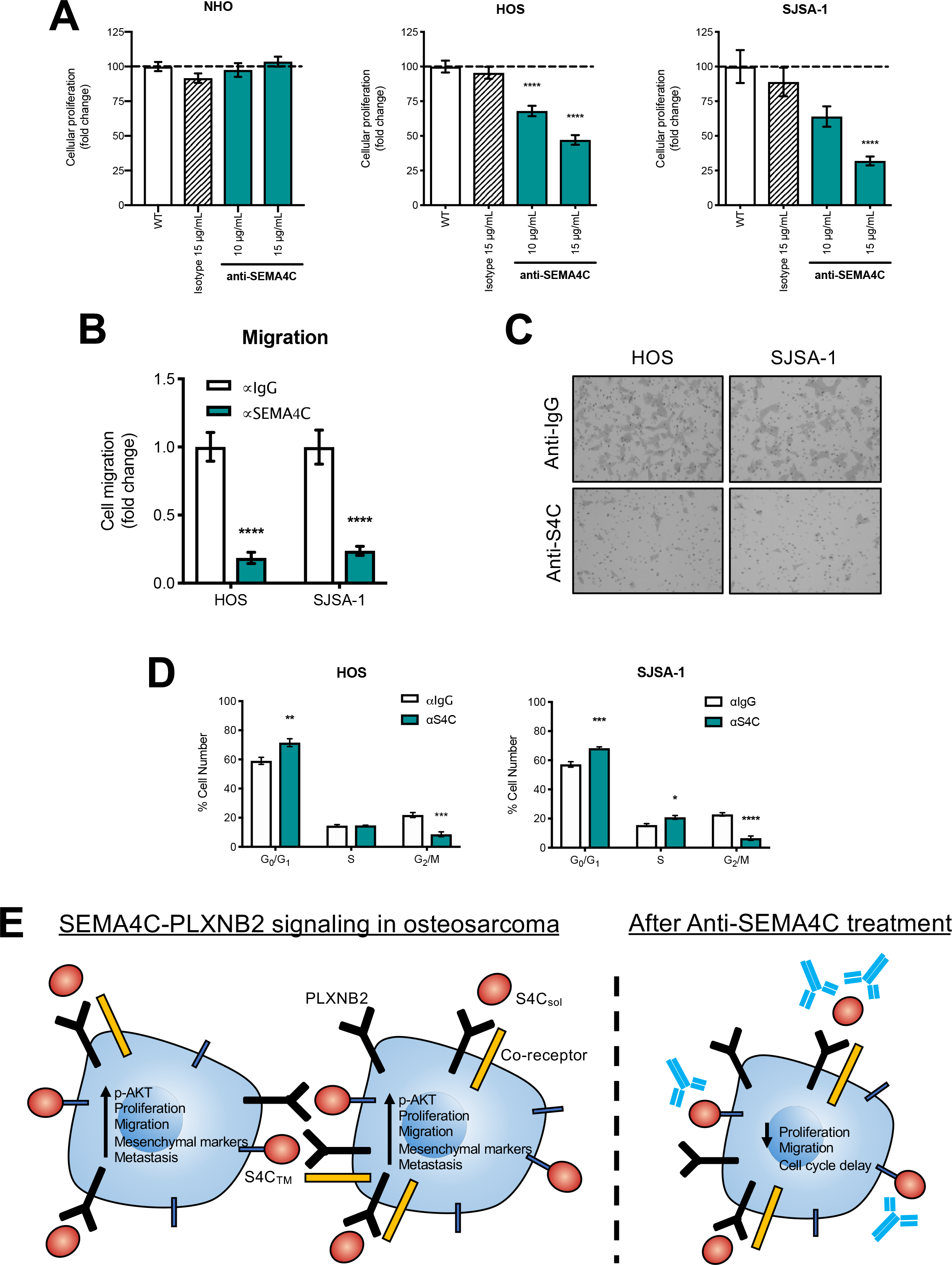
Anti-SEMA4C monoclonal antibody blockade is effective *in vitro*. **A.** SEMA4C antibody blockade reduces cellular growth following a 48-hour incubation in osteosarcoma cell lines only. Data as fold change ± SEM (n = 18); \*\*\*\**p* < 0.0001; One-way ANOVA. **B.** Antibody blockade reduces cellular migration. Data shown as fold change ± SEM (n = 12); \*\*\*\**p* < 0.0001; Student’s T-test. **C.** Representative images of migration in isotype control IgG and anti-SEMA4C treated lines. **D.** Anti-SEMA4C treatment induces G1 cell cycle delay. Data shown as number of cells ± SEM (n = 3); \**p* < 0.05, \*\**p* < 0.01, \*\*\**p* < 0.001, \*\*\*\**p* < 0.0001; Two-way ANOVA. **E.** Model of SEMA4C function. SEMA4C promotes downstream activation of AKT signaling which ultimately leads to upregulation of mesenchymal genes, promotion of cellular migration, proliferation, tumor growth, and metastasis. Monoclonal antibody blockade can effectively inhibit these downstream events and induce cell cycle delay. S4C_TM_ = transmembrane SEMA4C, S4C_sol_ = soluble SEMA4C.

## 4. Discussion

Our studies provide several new insights into the functions of the semaphorin 4C (SEMA4C) signaling pathway promoting osteosarcoma progression and metastasis. SEMA4C has been shown to be highly expressed relative to control tissues and cell lines. Its heightened expression and signaling through its cognate receptor PLXNB2 correlates to patient outcomes in some solid cancers [30]. Our results demonstrate high level cytoplasmic/membraneous SEMA4C expression in malignant primary patient tissues which suggests SEMA4C is targetable on the cell surface. Activation of PLXNB2 signaling by its ligand SEMA4C is likely to occur via autocrine mechanisms in co-expressing tumor cells and via paracrine signaling when PLXNB2 is stimulated from SEMA4C-producing cells in the microenvironment. Our study demonstrates that both normal osteoblasts and osteosarcoma cell lines indeed express both soluble SEMA4C and cognate receptor PLXNB2, but interestingly membrane-bound SEMA4C was only found to be detectable in osteosarcoma cell lines. This data supports both autocrine and paracrine capabilities, but at an exaggerated level similar to previous findings in tumor-associated lymphatic endothelial cells and breast cancer [14, 27]. These elevated levels suggest that osteosarcoma cells are also genetically and phenotypically distinct from that of their normal counterparts.

Accumulating evidence suggests that SEMA4C-PLXNB2 interactions can promote an oncogenic signaling axis. Recent reports from Gurrapu et al. and Le et al. established that SEMA4C-PLXNB2 signaling promotes cancer cell proliferation, migration, and tumorigenesis in breast and glioma tumor cells respectively [14, 16]. In our study, we demonstrated that SEMA4C directs both 2D and 3D growth *in vitro* as well as modulates invasive cell motility and adhesion in osteosarcoma. These phenotypes were associated with large perturbations in activated AKT signaling. In particular, genetic silencing of SEMA4C induced G1 cell cycle delay, which has been well-established and linked with changes in PI3K pathway signaling [31]. These *in vitro* findings were further substantiated by our mouse studies. We injected two SEMA4C knockdown cell lines into mice using a highly relevant orthotopic mouse model of osteosarcoma [24, 25]. SEMA4C knockdown tumor cells exhibited reduced tumor growth, micrometastatic nodule formation, and area. These data indicate a diverse role for SEMA4C in osteosarcoma growth, metastasis, and maintenance.

Among the family of SEMA4 members, several recent reports have demonstrated that SEMA4C also regulates EMT [32, 33]. While this phenomenon has been well-established in cancers of epithelial origin [34], the precise and novel roles EMT factors play in a characteristically mesenchymal cancer such as osteosarcoma, remain to be determined. In agreement with other reports in controlling EMT [32, 33], our results suggest high level SEMA4C expression promotes invasive cell motility, is associated with mesenchymal marker expression, and can be reversed through genetic disruption. Excitingly, TWIST1 was significantly altered in both of our SEMA4C gain-of-function (GOF) and loss-of-function (LOF) studies. Research from Yin and colleagues suggests TWIST1 is associated with poor prognoses in osteosarcoma and can be used as a prognostic indicator of metastatic potency in patients [35]. Moreover, in a study of 206 unique bone tumors, TWIST1 expression was one of three markers in a panel that afforded the most sensitive and specific diagnostic utility among varied bone tumor types [36]. TWIST1 is also essential for tumor initiation and maintaining a mesenchymal state in synovial sarcomas [37]. Reactivation of TWIST1 can also promote metastasis in other sarcomas such as Ewing’s sarcoma [38]. The continued demonstration of the plasticity of TWIST1 and other mesenchymal markers in maintaining phenotypes associated with many aspects of osteosarcomagenesis, progression, and metastasis has led to the central belief of skeletal cancer stem cells [39, 40], of which have been just recently identified [41]. High level expression of SEMA4C may help facilitate a hyper-mesenchymal state in osteosarcoma and/or even allow a plasticity that contributes to many of the phenotypes our study illustrates, including wound healing [42], cellular adhesion [43], tumor growth, and metastasis [44].

The findings of our work may be highly relevant to the clinical setting. To date, metastases to the lungs remains the number one cause of osteosarcoma-related death [45]. Our monoclonal antibody blockade studies support the concept of targeting SEMA4C therapeutically. Targeted blockade of SEMA4C-induced signaling appreciably slowed tumor cell proliferation and migration *in vitro*. These phenotypic changes were again accompanied by G1 cell cycle delay. Together, these data suggest a rationale for SEMA4C blockade as a potential novel treatment option for patients with metastatic osteosarcoma. Recent studies on SEMA4D, a type IV semaphorin member that can also signal through PLXNB2 [46], suggests high level expression restricts tumoricidal immune cells from entering the tumor microenvironment and blunts their activity, however, monoclonal antibody neutralization by an anti-Sema4d antibody (murine: mAb67-2, human: VX15/2503; Vaccinex, Inc.) could restore these deficits in combination with anti-Ctla-4 or anti-Pd-1 checkpoint blockade [47–49]. Likewise, antibody targeting of SEMA4C could be highly advantageous for these same reasons and others we posit here. This could allow expansion of the poor portfolio of therapies available to metastatic osteosarcoma patients and may also be applied to other cancer types in which high SEMA4C expression is clinically relevant.

SEMA4C positive tumor cells are an attractive target therapeutically. These results suggest the possibility of SEMA4C as a novel therapeutic target for the treatment of incurable metastatic osteosarcoma.

## 5. Acknowledgements

The authors would like to thank Dr. Kyle Williams for helpful discussion throughout the study, Dr. Sterbs for her intoxicating enthusiasm during the preparation of this manuscript, Dr. Juan Abrahante for statistical advice, and the Clinical and Translational Science Institute (CSTI) Histology and Research Laboratory team members Dr. Colleen Forester and Lori Holm for tissue preparation and histology services. Author B.A.S. is supported by an NIH NIAMS T32 AR050938 Musculoskeletal Training Grant. Author E.J.P. is supported by an NIH NIAID T32 AI997313 Immunology Training Grant. This work was made possible through funding from the Sobiech Osteosarcoma Fund Award, Randy Shaver Cancer and Community Fund, University of Minnesota Foundation, Rein in Sarcoma Foundation, Aflac-AACR Career Development Award, and the Children’s Cancer Research Fund to author B.S.M. Portions of this work were conducted in the Minnesota Nano Center, which is supported by the National Science Foundation through the National Nano Coordinated Infrastructure Network (NNCI) under Award Number ECCS-1542202.

## 6. Authors’ contributions

<u>Conception and design:</u> B.A.S., D.A.L., B.S.M.

<u>Development and acquisition of data:</u> B.A.S, N.J.S., E.J.P. H.E.B., G.A.S., M.R.C., J.J.P., G.M.D., K.L.B., E.P.R.

<u>Analysis and interpretation:</u> B.A.S., N.J.S., D.J.O., D.K.W. J.B.M., D.A.L, B.S.M.

<u>Writing, review and revisions:</u> B.A.S, N.J.S., E.J.P. H.E.B., G.A.S., M.R.C., J.J.P., G.M.D., K.L.B., E.P.R., J.B.M., D.J.O., D.K.W., D.A.L., B.S.M

<u>Study oversight:</u> B.A.S., D.A.L., B.S.M

---

**Supp. Table 1.** Table of all antibodies used in manuscript.

| **Antigen** | **Source** | **Dilution** | **Catalog** | **Company** |
| --- | --- | --- | --- | --- |
| SEMA4C | Mouse | 1:250 | 136445 | Santa Cruz Biotechnology |
| PLXNB2 | Sheep | 1:500 | AF5329 | R & D Systems |
| p-AKT (Ser473) | Rabbit | 1:1000 | 4060 | Cell Signaling Technologies |
| AKT | Rabbit | 1:1000 | 4691 | Cell Signaling Technologies |
| B-ACTIN | Mouse | 1:5000 | 3700 | Cell Signaling Technologies |

**Supp. Table 2.**
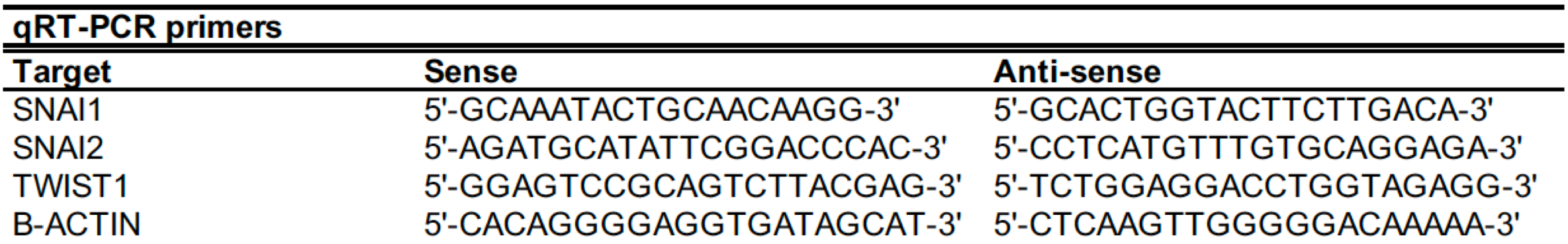
Table of primer sequences used in this manuscript.

**Supp. Fig. 1.**
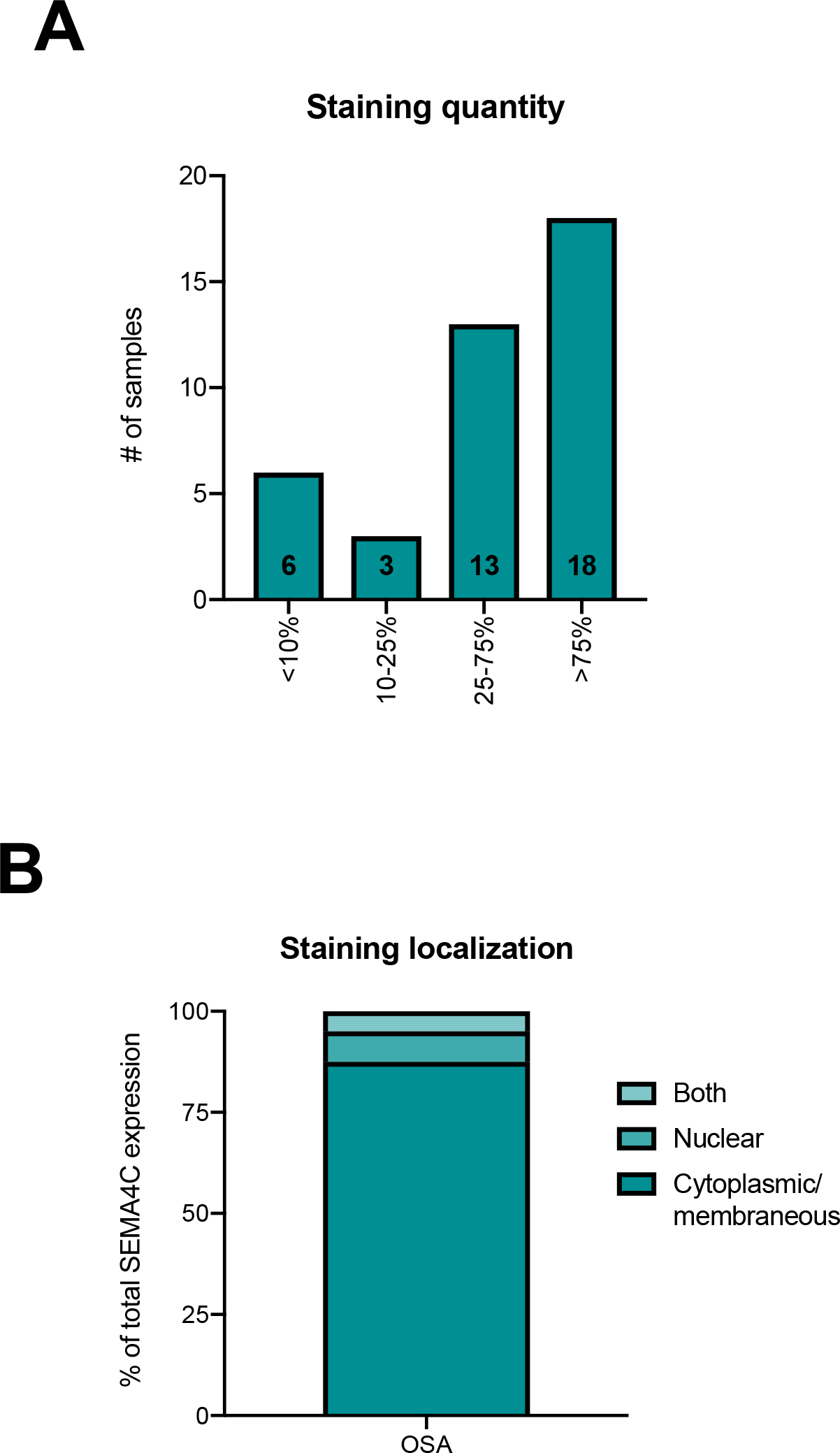
Increased SEMA4C expression is associated with osteosarcoma. **A.** Summary of the percent positive SEMA4C staining per section in a human osteosarcoma TMA. Data shown as number of samples with indicated percent staining in each of the four categories. **B.** Bar graph depicting percentage of SEMA4C staining localization (cytoplasmic/membraneous, nuclear, or both). SEMA4C staining was predominantly cytoplasmic/membraneous in sections.

**Supp. Fig. 2.**
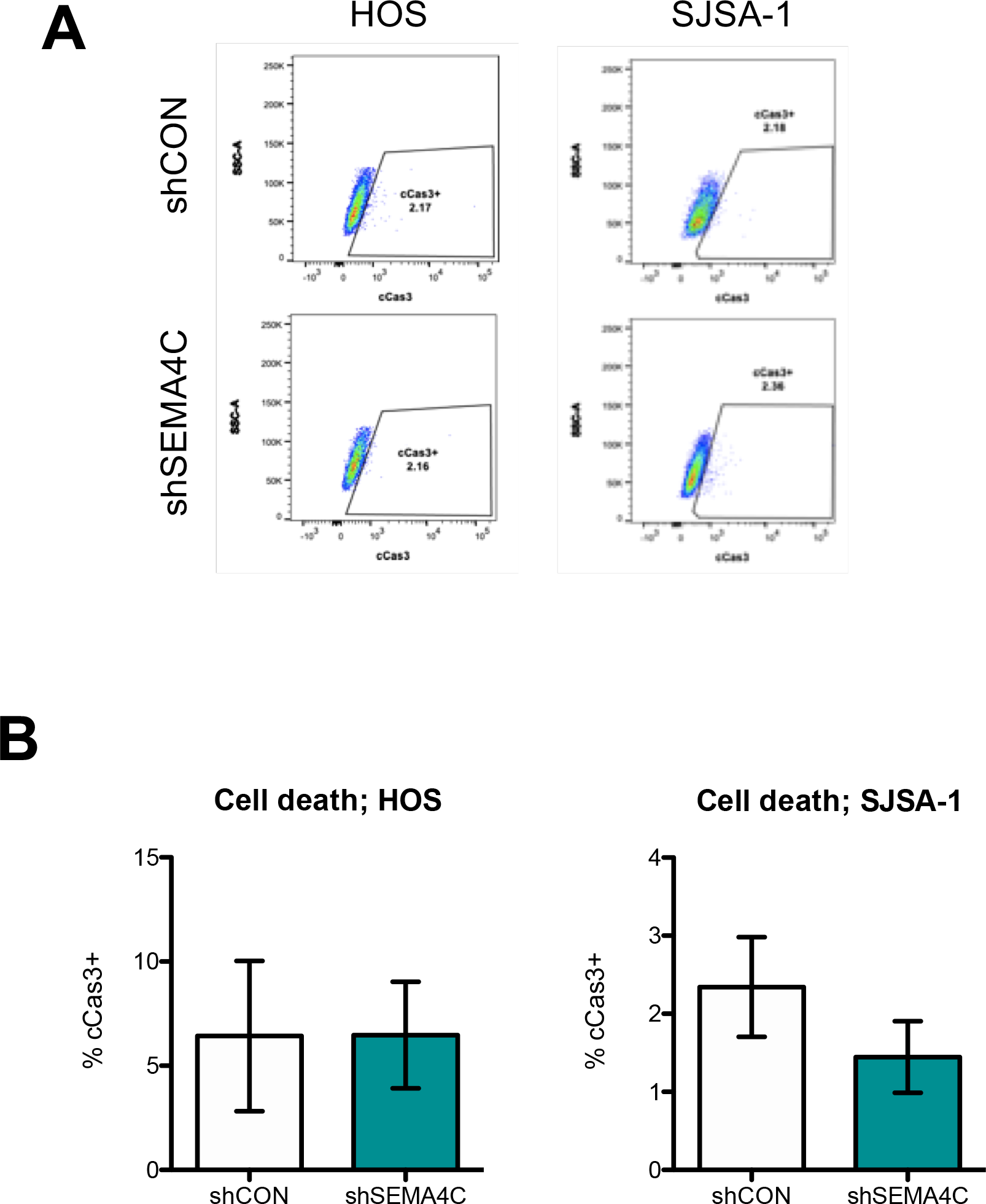
Silencing of SEMA4C does not induce apoptosis. **A.** Representative flow cytometry plots of cleaved CASP3 positivity (cCas3+) in SEMA4C knockdown cell lines. **B.** Quantification of flow cytometry plots. Data shown as mean area ± SEM (n = 3); *p* > 0.05; unpaired Student’s T-test.

**Supp. Fig. 3.**
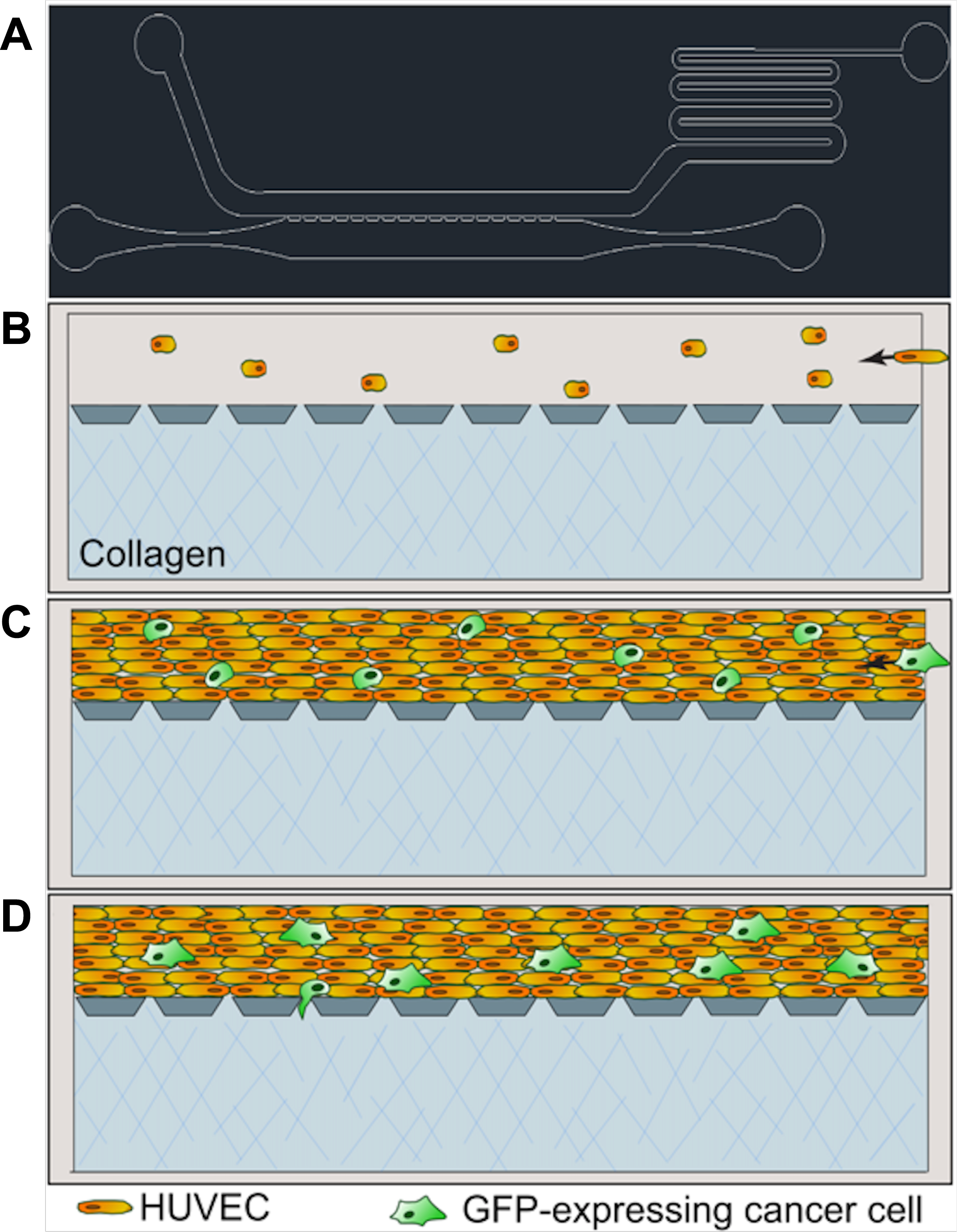
Description of 3D microfluidic chambers and representative images. **A.** An AutoCAD schematic of the entire device. **B.** Collagen (blue) is allowed to polymerize in the lower channel, then human umbilical vein endothelial cells (HUVECs, orange) are perfused through the adjacent channel. **C.** HUVECs are allowed to become confluent. Green fluorescent protein (GFP)-expressing cancer cells are perfused through the endothelial cell channel. **D.** Modified osteosarcoma cell lines adhere to the endothelium and may transmigrate and invade into the collagen.

**Supp. Fig. 4.**
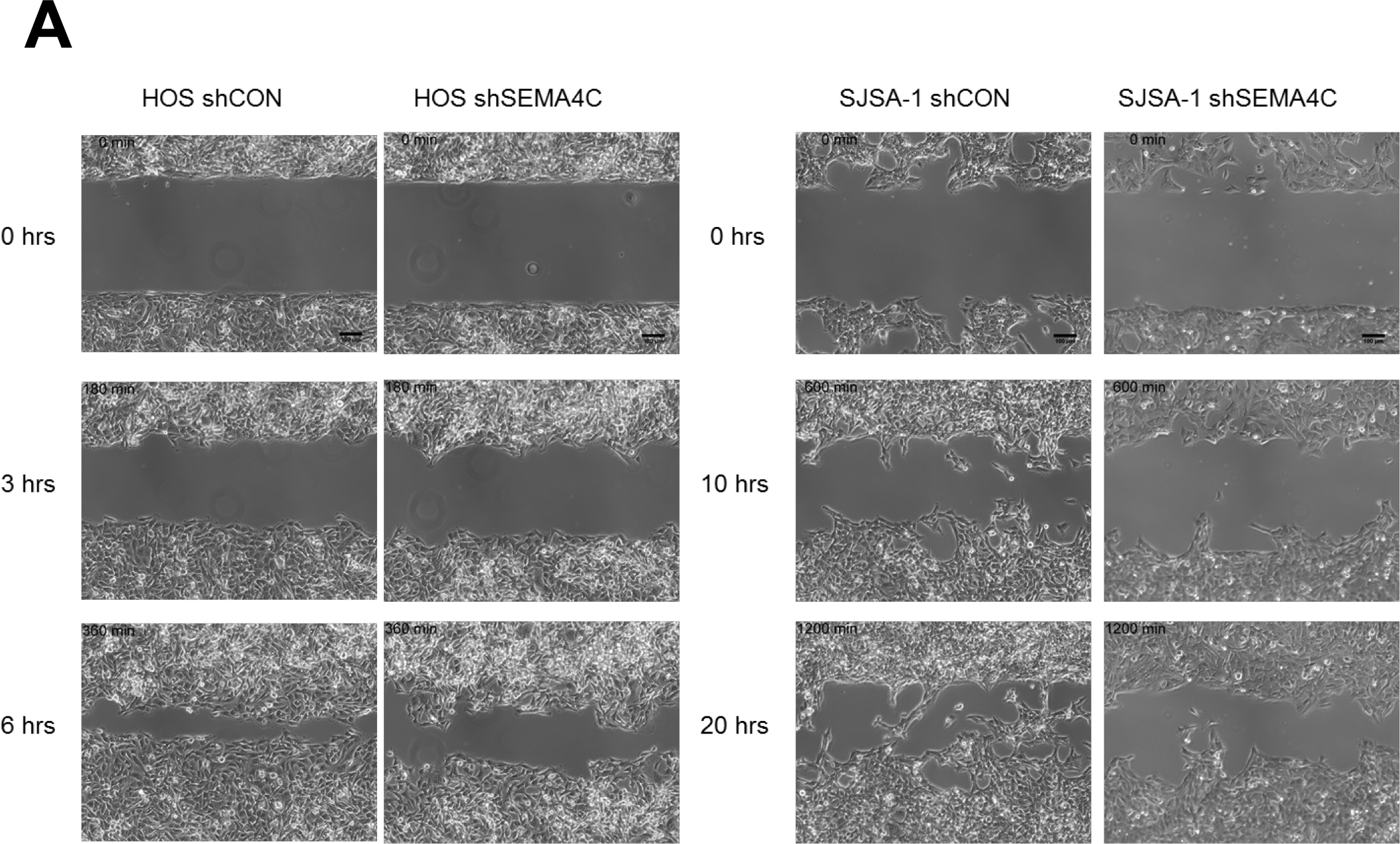
Representative illustration of wound closure assay analysis. **A.** Representative photo montages of phase contrast images of wound closure assays in cell lines ± SEMA4C deficiency at indicated time points.

